# The promoter regions of intellectual disability-associated genes are uniquely enriched in LTR sequences of the MER41 primate-specific endogenous retrovirus: an evolutionary connection between immunity and cognition

**DOI:** 10.1101/434209

**Authors:** Serge Nataf, Juan Uriagereka, Antonio Benitez-Burraco

## Abstract

Social behavior and neuronal connectivity in rodents have been shown to be shaped by the prototypical T lymphocyte-derived pro-inflammatory cytokine Interferon-gamma (IFNγ). It has also been demonstrated that STAT1 (Signal Transducer And Activator Of Transcription 1), a transcription factor (TF) crucially involved in the IFNγ pathway, binds consensus sequences that, in humans, are located with a high frequency in the LTRs (Long Terminal Repeats) of the MER41 family of primate-specific HERVs (Human Endogenous Retrovirus). However, the putative role of an IFNγ/STAT1/MER41 pathway in human cognition and/or behavior is still poorly documented. Here, we present evidence that the promoter regions of intellectual disability-associated genes are uniquely enriched in LTR sequences of the MER41 HERVs. This observation is specific to MER41 among more than 130 HERVs examined. Moreover, we have not found such a significant enrichment in the promoter regions of genes that associate with autism spectrum disorder (ASD) or schizophrenia. Interestingly, ID-associated genes exhibit promoter-localized MER41 LTRs that harbor TF binding sites (TFBSs) for not only STAT1 but also other immune TFs such as, in particular, NFKB1 (Nuclear Factor Kappa B Subunit 1) and STAT3 (Signal Transducer And Activator Of Transcription 3). Moreover, IL-6 (Interleukin 6) rather than IFNγ, is identified as the main candidate cytokine regulating such an immune/MER41/cognition pathway. Of note, functionally-relevant differences between humans and chimpanzees are observed regarding the 3 main components of this pathway: i) the protein sequences of immunes TFs binding MER41 LTRs, ii) the insertion sites of MER41 LTRs in the promoter regions of ID-associated genes and iii) the protein sequences of the targeted ID-associated genes. Finally, a survey of the human proteome has allowed us to map a protein-protein network which links the identified immune/MER41/cognition pathway to FOXP2 (Forkhead Box P2), a key TF involved in the emergence of human speech. Together, these results suggest that the stepped self-domestication of MER41 in the genomes of primates could have been a driver of cognitive evolution. Our data further indicate that non-inherited forms of ID might result from alterations of the immune/MER41/cognition pathway induced notably by the untimely or quantitatively inappropriate exposure of human neurons to IL-6.

## INTRODUCTION

Interferon gamma (IFNγ), the prototypical T Helper 1 (TH1) cytokine, is a T-cell derived pro-inflammatory molecule exerting several effects on innate immune cells and others on non-immune cells including neurons (Litteljohn et al., 2014). Specifically, IFNγ was recently shown to be a social behavior regulator and to shape neuronal connectivity in rodents (Filiano et al., 2016). Irrespective of cell type, binding of IFNγ to its receptor induces the transcriptional regulation of target genes via the recognition of promoter consensus sequences by the transcription factor STAT1 (Signal Transducer And Activator Of Transcription 1) (Green et al., 2017; Ramana et al., 2002). In humans, an important share of the IFNγ/STAT1 pro-inflammatory pathway is mediated by the binding of STAT1 to consensus sequences localized in the LTRs (Long Terminal Repeats) of the MER41 family of HERVs (Human Endogenous Retroviruses) (Chuong et al., 2016). Thus, in primates only, MER41 sequences located in the promoter regions of immune genes, serve as IFNγ-inducible enhancers that are indispensible for a full IFNγ-mediated immune response (Chuong et al., 2016). MER41 integrated into the genome of a primate ancestor 45-60 million years ago and a total of 7,190 LTR elements belonging to 6 subfamilies (MER41A-MER41G) are detectable in the modern human genome (Chuong et al., 2016). That being said, the hypothesis of an IFNγ/STAT1/MER41 pathway shaping social behavior and/or cognition in humans (Filiano et al., 2016) is not yet supported by experimental data. Addressing this issue is of importance as it could provide additional evidence on whether and how the immune system may translate environmental cues (including, possibly, cultural cues) into genomic regulatory pathways shaping behavior and/or cognition. Obviously, multiple research fields are concerned, from cognitive evolution to psychiatric disorders. As a first step, any human candidate gene(s) regulated by such a pathway need(s) to be identified. Additionally, it is important to determine if STAT1 is the sole immune TF that potentially regulate the transcription of behavior- or cognition-associated genes via MER41 LTRs in humans. Finally, it is worth considering whether the stepped integration of HERVs into the human genome, and more generally any evolutionary change dictated by infectious events, could be related to key cognitive specificities experienced by hominins. Indeed, it was recently hypothesized that the horizontal transfer of genetic material by viral and non-viral vectors might have prompted the emergence of language in the human species (Benítez-Burraco and Uriagereka, 2016).

To address these issues, we followed a bioinformatics workflow relying on the use of two recently-generated web tools allowing a survey of HERV sequences and their associated transcription factor binding sites (TFBSs) in the entire human genome. Using this approach, we found that the promoter regions of genes causatively linked to genetically-determined intellectual disability (ID) are highly significantly enriched in LTR sequences of the MER41 family. Such an enrichment was unique to both MER41, as compared to more than 130 explored HERV, and to ID-associated genes, as compared to lists of genes associated with autism spectrum disorder (ASD) or schizophrenia. The MER41 LTRs that localize in the promoter regions of ID-associated genes harbor binding sites recognized by canonical immune TFs including STAT1, STAT3 and NFKB1. From these data, we performed phylogenetic comparisons between humans and chimpanzees regarding: i) MER41 LTR insertion sites in the promoter regions of candidate ID-associated genes, ii) protein sequences of the immune-related TFs binding MER41 LTRs in such promoter regions and iii) protein sequences of the targeted ID-associated genes. This was so as to infer putative consequences for the human-specific expression pattern of ID-associated genes, which might ultimately account for some of the cognitive differences between species. Our results indicate a possible evolutionary advantage to humans in the immune/MER41/cognition pathway. Finally, since FOXP2 is currently known as a relevant TF regulating aspects of brain development and functions which are important for the execution of speech-related motor programs (Konopka et al., 2009; Oswald et al., 2017; Spiteri et al., 2007; Vernes et al., 2007, 2011), we searched genomics and proteomics databases to map a putative functional interactome linking FOXP2 and the immune TFs binding MER41 LTRs. This way, we were able to unravel a HERV-driven evolutionary-determined connection between cognition and immunity with a potential impact on language evolution and the pathophysiology of ID.

## MATERIALS AND METHODS

A scheme summarizing the workflow followed in the present work is shown in Figure 1.

**Figure 1:**
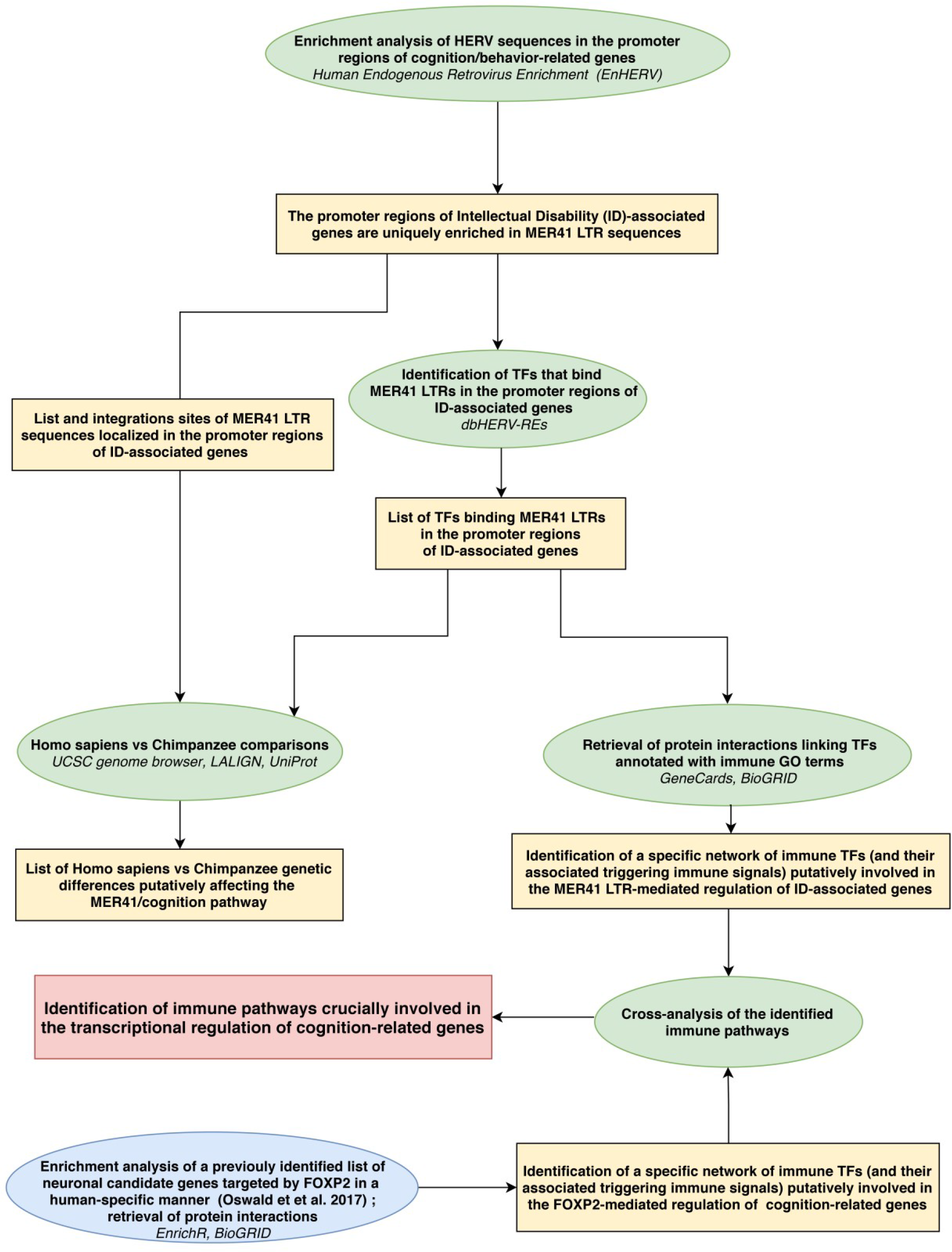
Workflow of the sudy. Rectangles (yellow or red) frame the main results obtained following each of the analytical steps briefly described in ellipse shapes (green or blue). Terms in italics correspond to the name of the bioinformatics tools used for each analytical step. LTR: long terminal repeats; ID: intellectual disability; TFs: transcription factors.

All the bioinformatics analyses were performed at least 3 times between December 2017 and February 2019. Bioinformatic tools and corresponding tasks performed in this study are described below.
- <u>The EnHERV database and web tool (</u>Tongyoo et al., 2017): identifying human genes harboring MER41 LTR sequence(s) in the promoter region located 2 KB upstream the TSS; only solo LTRs oriented in the sense direction relative to the gene orientation were taken into account.
- <u>The db-HERV-REs database and web tool (</u>Ito et al., 2017): identifying experimentally-demonstrated TFBSs in HERV LTRs. The db-HERV-REs database has been generated by the re-analysis of 519 ChIP-Seq datasets provided by the ENCODE (Davis et al., 2018; ENCODE Project Consortium., 2012; ENCODE Project Consortium, 2004) and Roadmap (Roadmap Epigenomics Consortium et al., 2015) consortia.
- <u>The enrichment web platform Enrichr (</u>Kuleshov et al., 2016): performing enrichments analyses on queried lists of genes. The Enrichr website allows surveying simultaneously 132 libraries gathering 245,575 terms and their associated lists of genes or proteins. Enrichment analysis tools provided by the Enrichr bioinformatics platform provides adjusted P-values computed from the Fisher’s exact test, Z-scores assessing deviation from an expected randomly obtained rank, and combined scores computed from the Z-scores and adjusted P-values obtained with the Fisher exact test. We essentially focused our analysis on the well-recognized “GO term biological process” library (Ashburner et al., 2000; The Gene Ontology Consortium, 2019) and on 3 ontology libraries based exclusively on text-mining: i) the “Jensen TISSUES” library (Santos et al., 2015), to determine whether a list of genes is significantly associated with a specific tissue or cell type, ii) the “Jensen COMPARTMENTS” library (Binder et al., 2014), to determine whether a list of genes is significantly associated with a specific cellular compartment or macromolecular complex and iii) the “Jensen DISEASES” library (Pletscher-Frankild et al., 2015), to determine whether a list of genes is significantly associated with a specific disease.
- <u>The UCSC genome browser (</u>Rosenbloom et al., 2015): retrieving the sequences of MER41 LTRs and their precise localization in the promoter region of ID-associated genes in the human genome (Human genome assembly GRCh38/hg38) and in the Pan troglodytes genome (Chimpanzee genome assembly CSAC 2.1.4/panTro4).
- <u>The Swiss Institute of Bioinformatics (SIB) sequence alignment web tool LALIGN (SIB Swiss Institute of Bioinformatics Members, 2016)</u>: performing sequence comparisons between human and chimpanzee MER41 LTRs located in the promoter region of ID-associated genes. For each of these genes we checked the presence, nature and precise localization of MER41 LTR sequences in the promoter region.
- <u>The UniProt database of protein sequence and functional information (The UniProt Consortium, 2018):</u> performing protein sequence alignments between Homo sapiens and Pan troglodytes for: i) TFs binding MER 41 LTRs in the promoter regions of ID-associated genes and ii) ID-associated genes harboring MER 41 LTRs in their promoter regions
- <u>The Brain RNA-Seq database (</u>Zhang et al., 2016): exploring mRNA expression profiles obtained by RNA-Seq analyses in primary cultures of human neurons, astrocytes or macrophages/microglia.
- <u>The TISSUES database (</u>Palasca et al., 2018): determining, for a given gene, which tissues harbor the highest levels of expression across a large range of normal human tissues. This database compiles results from 4 large expression atlases generated by pan-genomic and/or pan-proteomic analyses of normal human tissues (Clark et al., 2007; Fagerberg et al., 2014; Krupp et al., 2012; Su et al., 2004).

## RESULTS

### 1 The promoter regions of ID-associated genes are uniquely enriched in MER41 LTR sequences

We queried the EnHERV database and web tool (Tongyoo et al., 2017) to determine whether candidate lists of cognition/behavior-related genes were enriched in genes harboring promoter-localized HERV LTRs (more precisely: sense-oriented solo HERV LTR sequence(s) localized in the promoter region located 2 KB upstream the TSS). We performed such an analysis successively for the 133 families of HERV that can be mined on the EnHERV website. Three lists of cognition/behavior-related genes were assessed (data supplement 1): i) a list of high confidence ASD susceptibility genes established by the SFARI consortium (Abrahams et al., 2013) and based on expert-operated manual curation of the literature, ii) a recently-established list of putative schizophrenia-causing genes inferred from the integrative analyses of genome wide association studies (Ma et al., 2018), iii) a list of genes for which mutations or deletions are considered as causative of intellectual disability based on a manual curation of the literature (Kochinke et al., 2016). As indicated in the original paper describing the EnHERV web tool (Tongyoo et al., 2017), results were considered as statistically significant when both following criteria were fulfilled: a Fisher exact test P-values < 0.001 and an odds ratio > 1. Using this approach, we found that the promoter regions of ID-associated genes were highly significantly enriched in MER41 LTRs (P-value = 0.0004; odds ratio = 4.28). Results were not significant for any of the other 132 HERV families that can be mined on the EnHERV website nor for the promoter regions of ASD- or schizophrenia-associated genes. Further supporting the specificity of our findings, when analyzing the 22 lists of non-CNS related genes provided as training lists by the EnHERV server, we did not find any significant enrichment in genes with promoter-localized MER41 LTR sequences. To confirm our findings, we retrieved from the EnHERV website the whole list of coding genes which, in humans, harbor a sense-oriented promoter-localized MER41 LTR sequence. On this list of 79 genes (data supplement 2), we then performed enrichment analyses using the Enrichr website as described in the Materials & Methods section. We found no statistically significant enrichments with regard to “biological process” GO terms, tissue-specific expression or sub-cellular localization of gene products. However, text-mining enrichment analysis unraveled a significant enrichment in genes associated with the term “Intellectual disability” (Table 1).

**Table 1.**
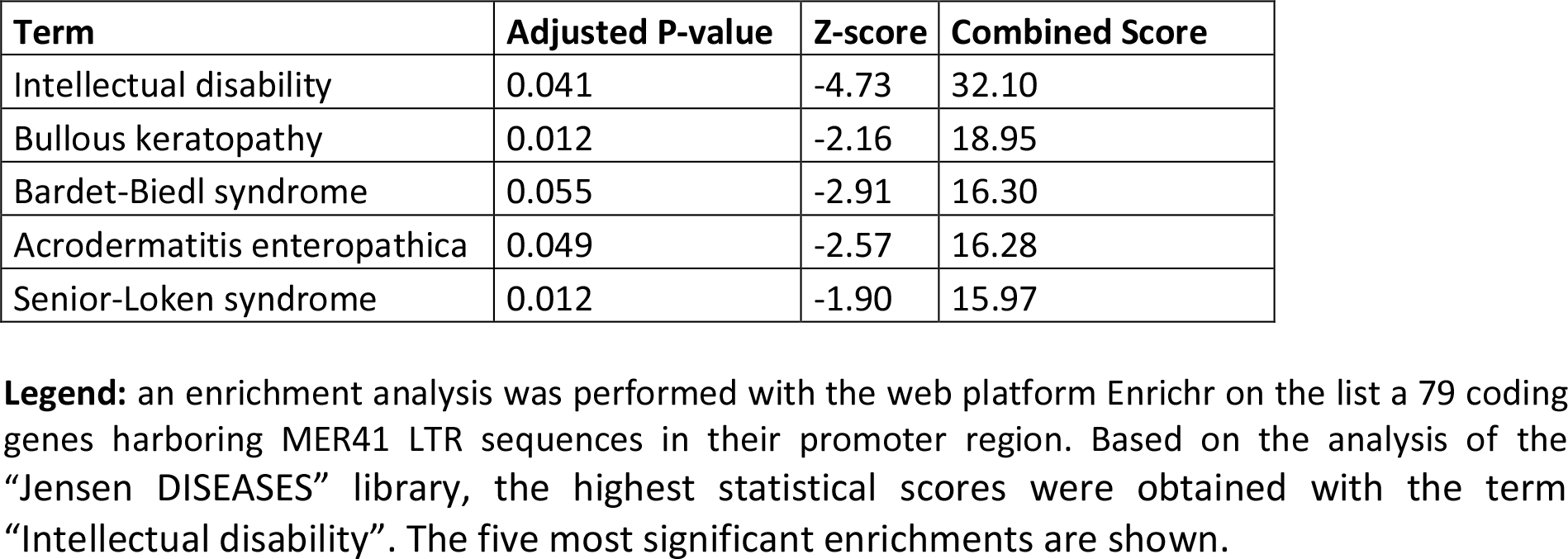

Since enrichment analysis based on text mining may be biased by the identification of a non-causative link between a given gene and the term “Intellectual disability”, we took into account only genes that had been identified as causative of ID (Kochinke et al., 2016). On this basis, out of 79 human genes harboring a MER41 LTR sequence in their promoter region, 9 had an established causative link with ID. The genes and associated genetic conditions are summarized in Table 2 and described below:

- *BBS10* (Bardet-Biedl syndrome 10): Bardet-Biedl syndrome 10 (vision loss, obesity, polydactily, kidney abnormalities and intellectual disability)
- *DEAF1* (DEAF1 transcription factor): Mental retardation, autosomal dominant 24 (intellectual disability and impairments in adaptive behavior)
- *AP1S1* (Adaptor Related Protein Complex 1 Subunit Sigma 1): MEDNIK syndrome (Mental retardation, enteropathy, deafness, peripheral neuropathy, ichthyosis and keratoderma)
- *ST3GAL5* (ST3 Beta-Galactoside Alpha-2,3-Sialyltransferase 5): Salt and pepper developmental regression syndrome (epilepsy, abnormal brain development and intellectual disability)
- *CDH15* (Cadherin 15): Mental retardation, autosomal dominant 3 (intellectual disability and impairments in adaptive behavior)
- *CEP290* (Centrosomal Protein 290): Bardet-Biedl syndrome 14 (vision loss, obesity, type 2 diabetes, hypercholesterolemia, polydactily, intellectual disability, impaired speech, delayed psychomotor development and behavioral alterations)
- *GAMT* (Guanidinoacetate N-Methyltransferase): Cerebral creatine deficiency syndrome 2 (epilepsy, intellectual disability and altered speech development)
- *DDHD2* (DDHD Domain Containing 2): Spastic paraplegia 54, autosomal recessive (delayed psychomotor development, intellectual disability and early-onset spasticity of the lower limbs)
- *GCSH* (Glycine Cleavage System Protein H): Glycine encephalopathy (hypotonia, delayed psychomotor development and epilepsy)

**Table 2.**
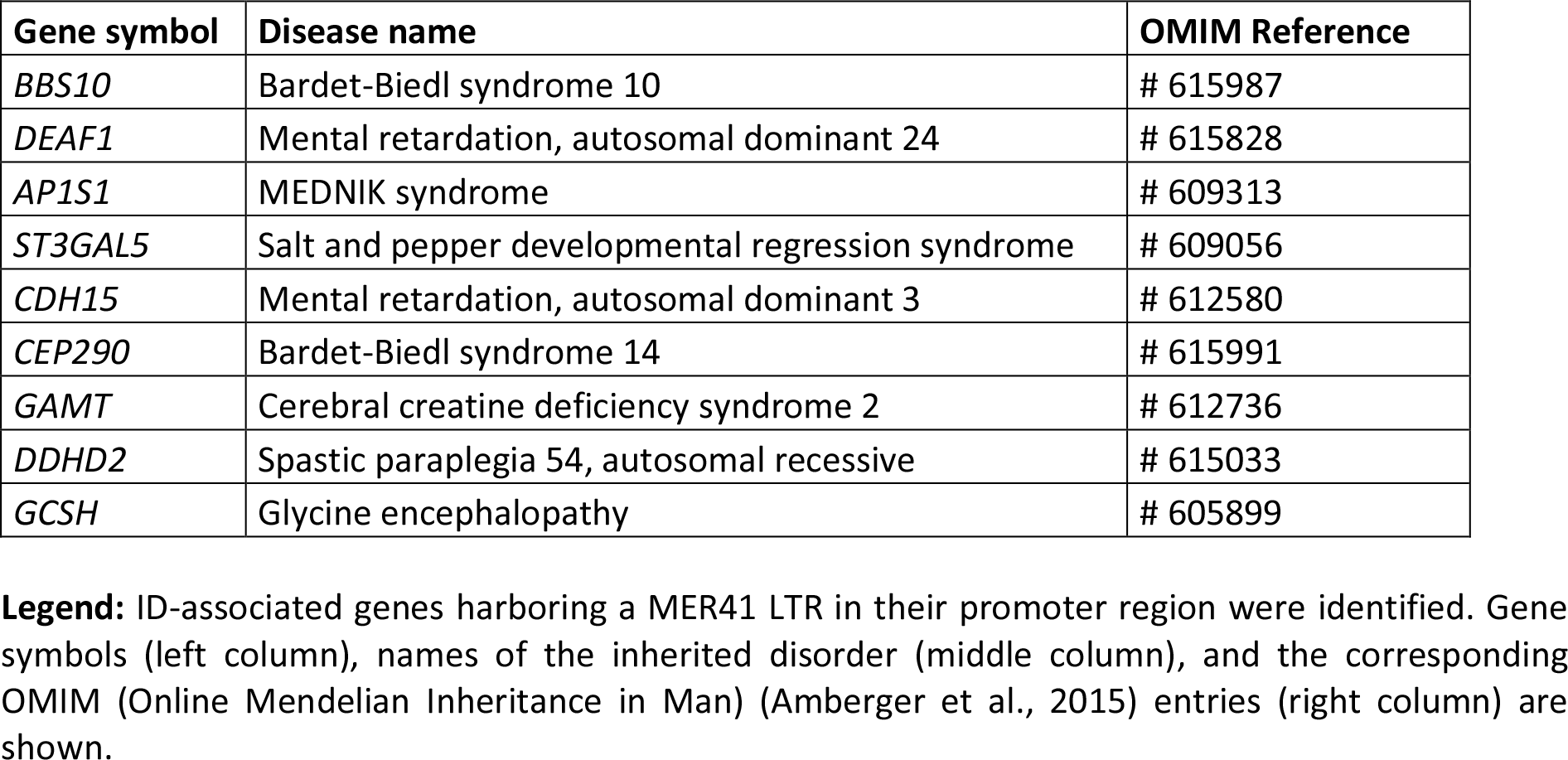

The “biological process” GO terms that annotate those 9 genes are shown in data supplement 3. Overall, these data point to a yet unrecognized potential link between promoter-localized MER41 LTRs and cognition.

### 2 LTRs from distinct members of the MER41 family of HERVs are inserted in the promoter regions of ID-associated genes

TFBSs in LTRs from the MER41 family (MER41 A-E and MER41G) have been shown to vary depending of the MER41 member considered (Chuong et al., 2016). Using the EnHERV database and web tool, we have identified several MER41 members for which LTRs can be demonstrated in the promoter regions of ID-associated genes. As shown in Table 3, only 3 ID-associated genes harbored a MER41B LTR in their promoter region: *CEP290*, *DDHD2* and *GCSH*. This indicates a potential transcriptional regulation of these 3 genes by the IFNγ/STAT1 pathway.

**Table 3.**
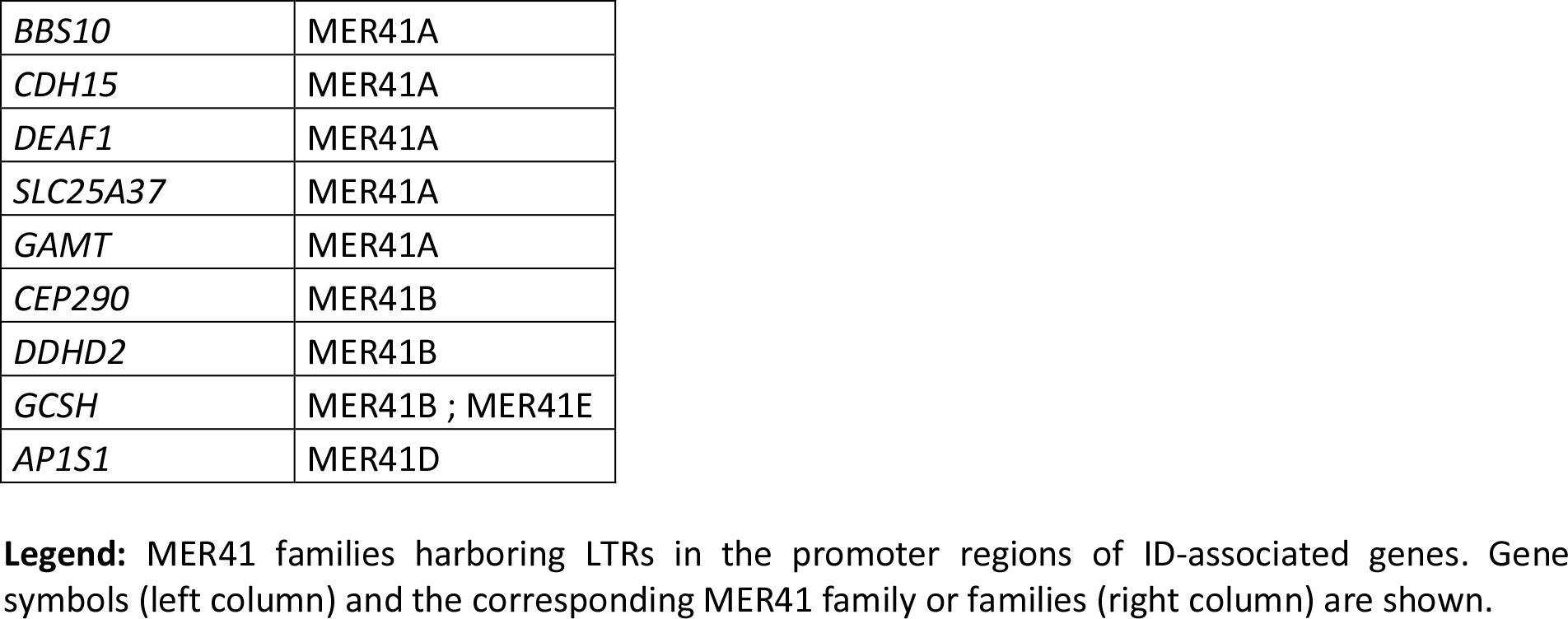

Interestingly, MER41 LTRs located in the promoter regions of ID-associated genes also include MER41A LTRs, which lack STAT1 binding sites (Chuong et al., 2016). This observation urged us to determine if other immune pathways (non IFNγ/STAT1-mediated) may regulate the transcription of ID-associated genes via MER41 LTRs. To this aim we used the HERV database and web tool “db-HERV-RE” (Ito et al., 2017), which allow the identification of experimentally-demonstrated TFBSs in HERV LTRs.

### 3 YY1 is the sole transcription factor harboring TFBSs in all the MER41 LTRs inserted in the promoter regions of ID-associated genes

Using the approach described above, we have identified 32 TFs that bind MER41 LTRs in the promoter regions of ID-associated genes (Table 4).

**Table 4.**
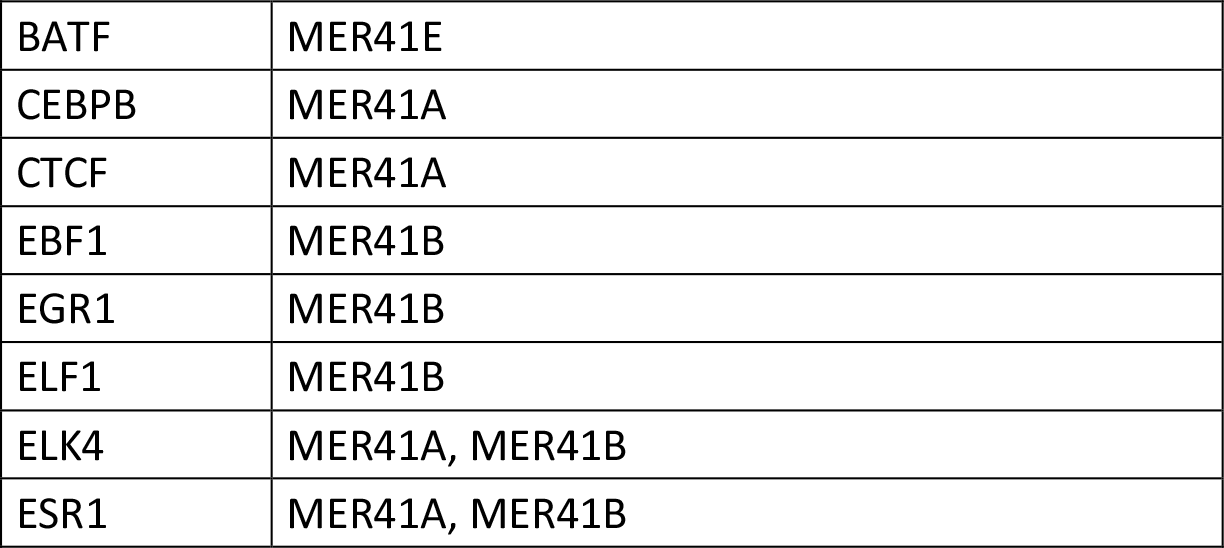
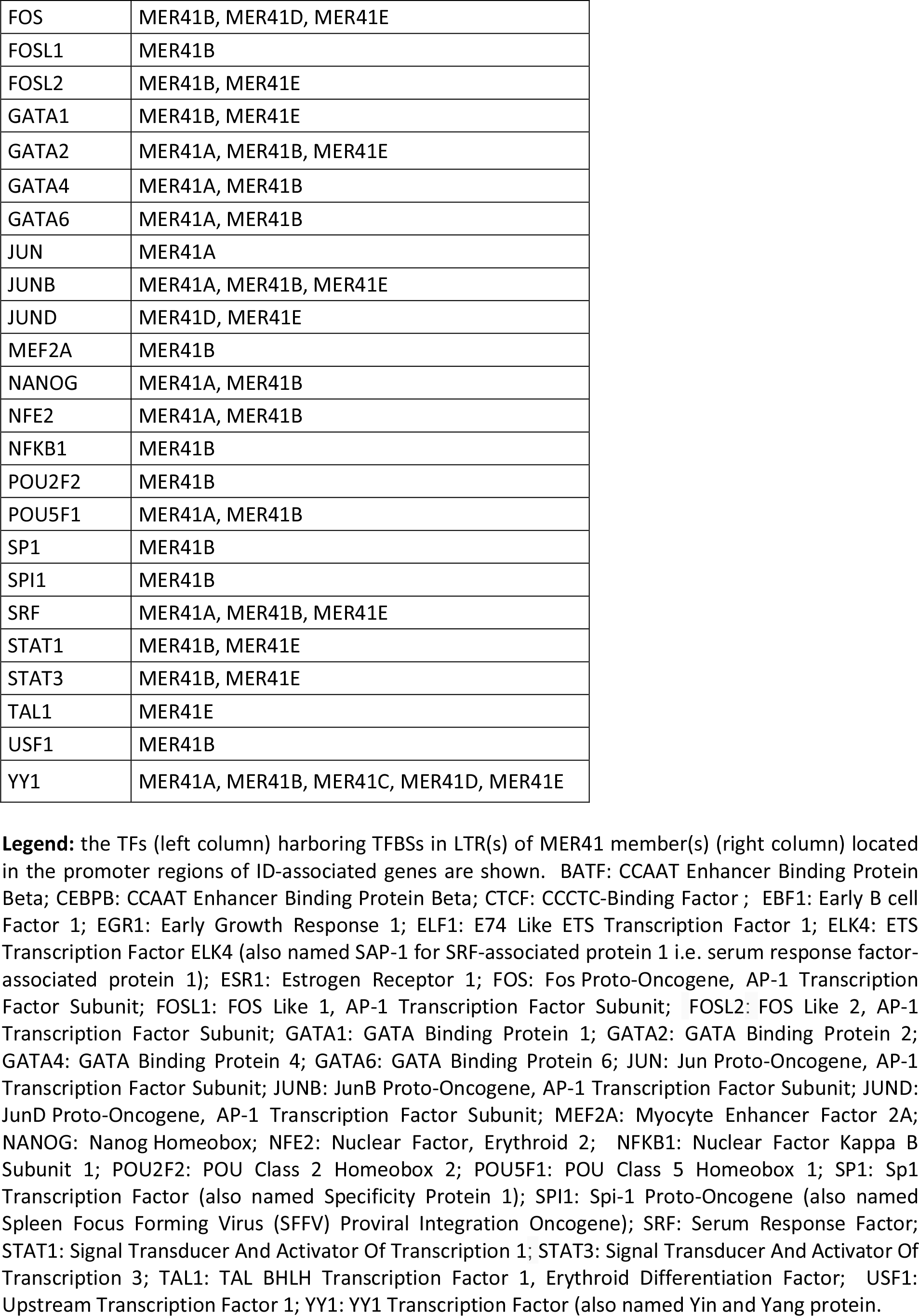

As expected, MER41B LTR comprises a STAT1 consensus sequence while MER41A LTR does not. Interestingly, an YY1 consensus sequence is present in the LTRs of all the MER41A-E members. It is worth noting that mutations/deletions in *YY1* are responsible for the Gabriele-De Vries syndrome, an autosomal dominant neurodevelopmental disorder characterized by intellectual disability, delayed psychomotor development and frequent autistic symptoms (Gabriele et al., 2017). Interestingly also, mutations in *CTCF*, another gene encoding a MER41 LTR-binding TF, are causally linked to “Mental retardation, autosomal dominant 21”, a developmental disorder characterized by significantly-below-average general intellectual functioning associated with impairments in adaptive behavior (Collins et al., 2016). Other inherited disorders associated to the above identified TF genes are summarized in Table 5 and notably include 3 groups of immune-related diseases induced by genetic alterations of the canonical immune TFs *STAT1*, *STAT3* and *NFKB1*, respectively.

**Table 5.**
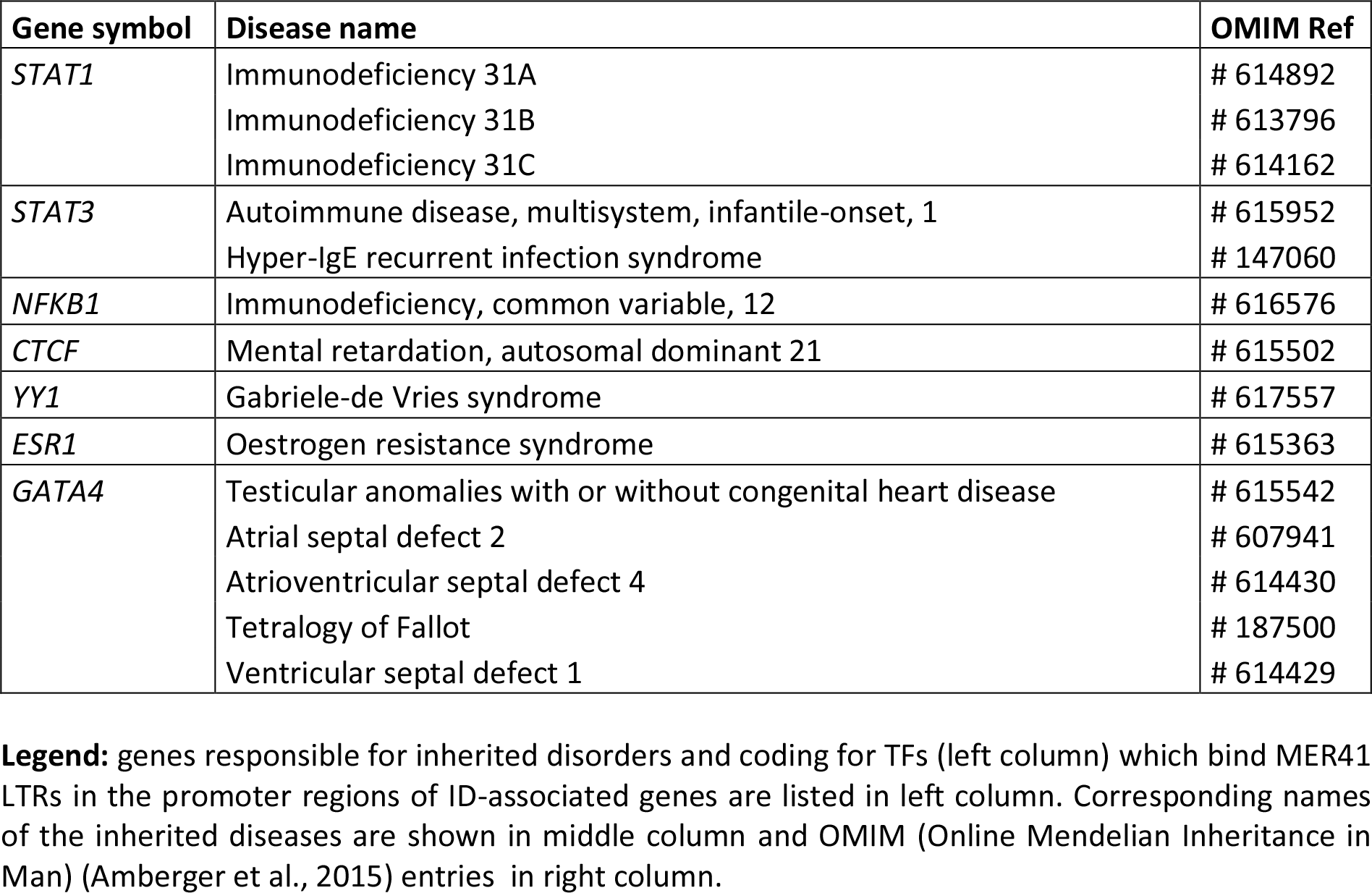

To summarize, besides STAT1, we have identified two canonical immune TFs, STAT3 and NFKB1, which bind specific MER41 LTRs in the promoter regions of ID-associated genes. Another TF, YY1, binds all MER41 LTRs in the promoter regions of ID-associated genes.

### 4 YY1 interact with a unique network of immune TFs that bind MER41 LTRs in the promoter regions of ID-associated genes

We also explored the BioGRID database of human protein interactions (Chatr-aryamontri et al., 2015) to determine whether STAT1, STAT3 and/or NFKB1 were reported to physically interact with each other and/or with YY1 and other TFs binding MER41 LTRs in the promoter regions of ID-associated genes. Interestingly, in the retrieved interaction network (Fig 2), we observed that YY1, via its interaction with NFKB1, is connected to a specific set of MER41 LTR-binding TFs that interact with STAT1, STAT3 and/or NFKB1. An analysis of the GO terms “Biological process” annotating each of these TFs indicates that besides STAT1, STAT3 and NFKB1, other members of this unique set of TFs exert immune functions (data supplement 4). This is notably the case for YY1. Moreover, some of such immune functions are linked to specific cytokines among which IL-1 and IL-6 are the most commonly shared in the retrieved GO terms (Fig 3). This result indicates that the prototypical pro-inflammatory molecules IL-1 and IL-6 are possibly involved in the transcriptional regulation of ID-associated genes displaying promoter-localized MER41 LTRs.

**Figure 2:**
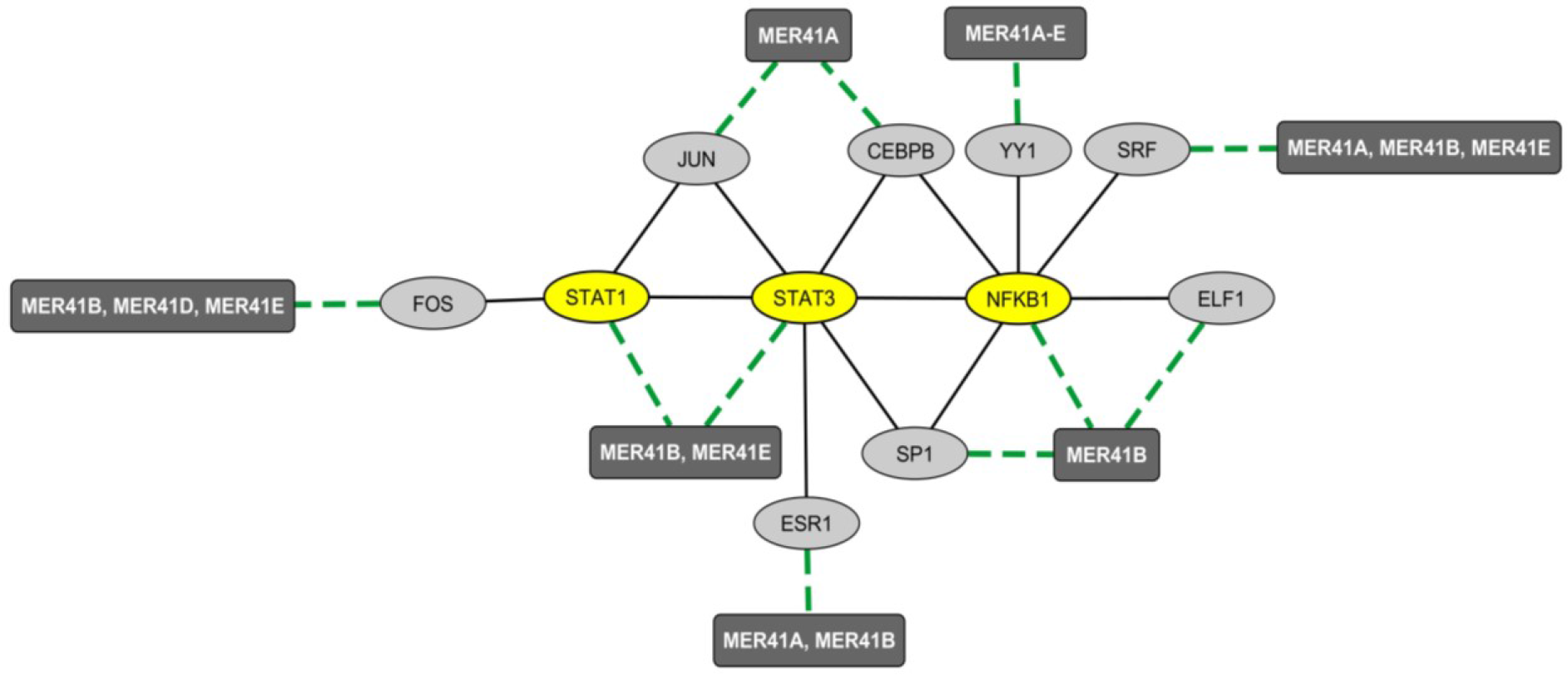
Mapping of the protein network formed by immune TFs that bind MER41 LTRs in the promoter regions of ID-associated genes. A survey of the human proteome was performed by querying the protein-protein interaction database BioGRID (Chatr-aryamontri et al., 2015). Ellipse shapes represent TFs and rectangles represent MER41 families displaying LTRs in the promoter regions of ID-associated genes. Black lines indicate protein-protein interactions. Green dashed lines indicate protein-DNA interactions between TFs and MER41 LTRs located in the promoter regions of ID-associated genes.

**Figure 3:**
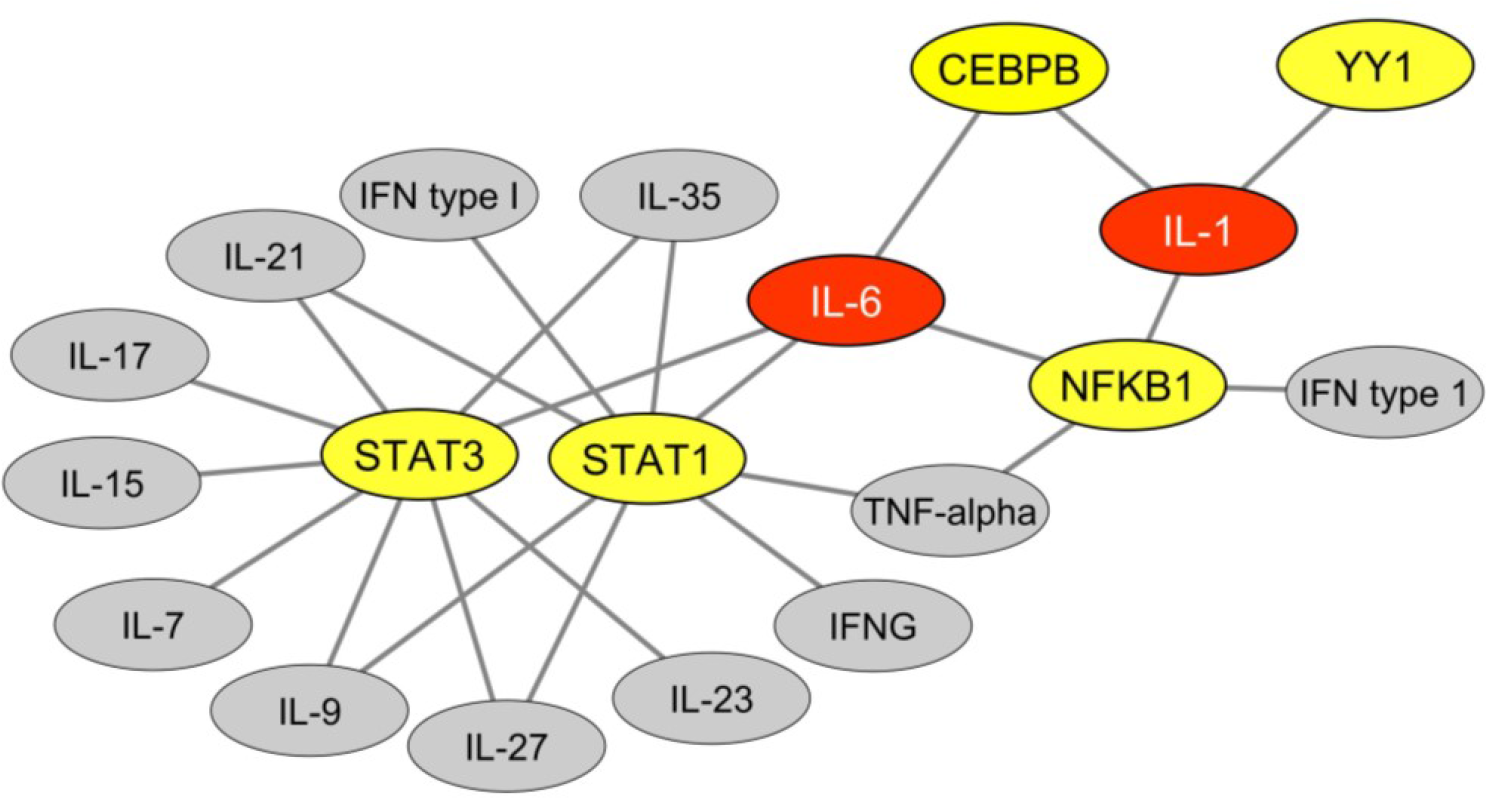
Cytokines functionally linked to TFs that bind MER41 LTRs in the promoter regions of ID-associated genes. An analysis was performed of the GO terms “biological process” that annotate each TFs which bind MER41 LTRs in the promoter regions of ID-associated genes. This allowed determining which cytokines are functionally linked with these TFs and potentially mediate an immune regulation of the MER41/cognition pathway. Such functional links are depicted as black lines. TFs are highlighted in yellow. Cytokines are highlighted in grey or, alternatively, in red for cytokines harboring 3 or more functional links with TFs. IFNG: interferon-gamma; IFN: interferon.

### 6 Chimpanzees vs Homo sapiens comparative analyses of the immune/MER41/cognition pathway

#### 6-1 MER41A-E LTR sequences and insertion sites in the promoter regions of cognition related (ID-associated) genes

As mentioned, MER41 HERVs integrated the genome of a primate ancestor 45-60 million years ago. The process of so-called “ERV domestication” (Dewannieux and Heidmann, 2013) relies on mechanisms that are not only species-specific, but may have partly shaped speciation (Johnson, 2015). Accordingly, in primates, the species-specific domestication of MER41 HERVs translates into the existence of species-specific differences regarding the insertion sites and/or sequences of integrated (fixed) LTRs. On this basis, we investigated whether the promoters of ID-associated genes harbored the same MER41A-E LTRs in human and chimpanzees (data supplement 5). Out of the 9 candidate genes examined we found that 5 exhibited, in both species, MER41 LTR sequences belonging to the same family and displaying 95 to 100% homology (data supplement 5). That being said, in 2 ID-associated genes MER41 LTR sequences were found at distances larger than 2 Kb from the TSS in chimps and, for 2 other genes (*CDH15* and *GCSH*), MER41 LTR sequences were absent, at least up to 10 Kb from the TSS in chimps. These results are indicative of differences that may prove functionally relevant with regard to the MER41 LTR-mediated transcriptional regulation of specific ID-associated genes. This remains to be experimentally explored. It is of note that, according to the classification provided by the Gene ontology (GO) consortium (The Gene Ontology Consortium, 2017), 3 of the genes displaying such promoter-localized differences are annotated with “Biological process” GO terms that may possibly render an account of distinctive features between chimpanzees and human CNS (data supplement 3). These include the terms “visual learning” and “locomotor behavior” for *DDHD2*, “hindbrain development” for *CEP290* and “glycine catabolic process” for GCSH (glycine being a major inhibitory neurotransmitter).

#### 6-2 Protein sequences of TFs binding MER41 LTRs in the promoter regions of cognition-related (ID-associated) genes

To further assess the potential role of MER41 LTRs in the species-specific transcriptional regulation of cognition-related (ID-associated) genes, the amino acid sequences of the 32 TFs that were reported to bind MER41A, MER41B, MER41D or MER41E LTRs in humans were then compared to those of their orthologs in chimpanzees. Using the UniProt-provided web tool “Align” we found that 15 out of the 32 examined TF sequences showed a 100% homology between humans and chimpanzees (data supplement 6). An additional set of 13 TFs exhibited 88.5% to 99.8% homology without any amino acid dissimilarities located in sequences of known or predicted functions. However, 4 TFs exhibited amino acid dissimilarities that are putatively functionally-relevant: YY1, ESR1, NANOG and SP1 (data supplement 6). Interestingly, the difference between the human and chimp YY1 sequences resides in the “Interaction with the SMAD1/SMAD4 complex” region which is essential to the modulating effects exerted by YY1 on the immunosuppressive TGF-beta pathway (Kurisaki et al., 2003).

#### 6-2 Protein sequences of cognition-related (ID-associated) genes harboring MER41 LTRs in their promoter regions

To further assess the existence of evolutionary-determined mechanisms that could have shaped the immune/MER41/cognition pathway in humans, we also performed comparisons between humans and chimpanzees regarding the amino acid sequences of the 9 proteins encoded by cognition-related (ID-associated) genes with promoter-localized MER41 LTRs (data supplement 6). In 6 out of these 9 proteins, we found amino acid dissimilarities that localized in sequences displaying known or predicted functions (The UniProt Consortium, 2018). Such functions notably include the interaction with identified protein partners, enzymatic activities, addressing toward specific subcellular compartments and interaction with DNA sequences (data supplement 6).

#### 6-4 Summary of humans vs chimpanzees comparisons

Overall, the observed differences are compatible with some evolutionary advantage to humans in the immune/MER41/cognition pathway. Such putative advantage may rely on: i) functional human specificities of TFs binding MER41 LTRs in the promoter regions of cognition-related (ID-associated) genes, ii) more proximal integration sites of MER41 LTRs in the promoter of specific cognition-related (ID-associated) genes, iii) functional human specificities of targeted cognition-related (ID-associated) genes harboring promoter-localized MER41 LTRs.

### 7 YY1 links FOXP2 to immune TFs binding MER41 LTRs in the promoter regions of cognition-related (ID-associated) genes

FOXP2, a TF abundantly expressed in cortical neurons, is involved in the emergence of human speech (Fisher and Scharff, 2009; Scharff and Petri, 2011; Vernes et al., 2011; Xu et al., 2018). Until recently, the transcriptional activity FOXP2 was thought to rely on the recognition of specific TFBs by FOXP2/FOXP2 homodimers or by FOXP1/FOXP2 or FOXP4/FOXP2 heterodimers (Sin et al., 2015; Wang et al., 2003). However, a recent work established a short list of TFs that bind FOXP2 and are likely to form heterodimers that regulate FOXP2 availability and/or DNA binding properties in neurons (Estruch et al., 2018). Interestingly, YY1 was identified as one of the 7 newly identified FOXP2-interacting TFs. We thus sought to determine whether YY1 could potentially represent a molecular link between FOXP2 and the immune/MER41/cognition pathway we identified. To this aim, we explored data obtained from a recent work attempting to identify a set of FOXP2 targets that are specific to human FOXP2 in neurons (Oswald et al., 2017). More precisely, data from this study were obtained by: i) a meta-analysis of previous works reporting on FOXP2 neuronal targets (based notably on Chip-Seq analyses of the neuronal cell line SH-SY5Y) (Enard et al., 2009; Hilliard et al., 2012; Konopka et al., 2009; Spiteri et al., 2007; Vernes et al., 2007, 2011) and ii) a comparison of neuronal genes that are targeted by human FOXP2 vs non-human primates orthologs of FOXP2 in the neuronal cell line SH-SY5Y (Oswald et al., 2017). A set of 40 candidate proteins encoded by FOXP2-targeted genes was identified. Protein interactors of these candidate targets were added in order to establish a final list of 80 proteins that are putatively regulated by FOXP2 in neurons in a human-specific manner. Interestingly, when performing enrichment analyses of the list of genes encoding such 80 proteins (data supplement 7), we found a highly significant enrichment in genes that are either associated with immune-related terms, as identified by text mining (“immune System”, “NFKB complex”, “arthritis” and others), or linked to immune biological processes according to the GO term classification (“cellular response to IL-21”, “cellular response to IL-2” and others). Of note, the list of genes putatively regulated by FOXP2 in a human-specific manner is significantly enriched in genes involved in “interleukin-6-mediated signaling pathway” (adjusted p-value: 0.0008) pointing again to IL-6 as a possible important player in the immune/MER41/cognition pathway. Finally, such a list of FOXP2 targets comprised STAT3 and several protein partners of STAT1, STAT3 and/or NFKB1. Integrating these data with the demonstrated interaction of FOXP2 with YY1 allows us to map a network of immune TFs that link FOXP2 to specific cognition-related (ID-associated) genes exhibiting promoter-localized MER41 LTRs (Fig 4).

**Figure 4:**
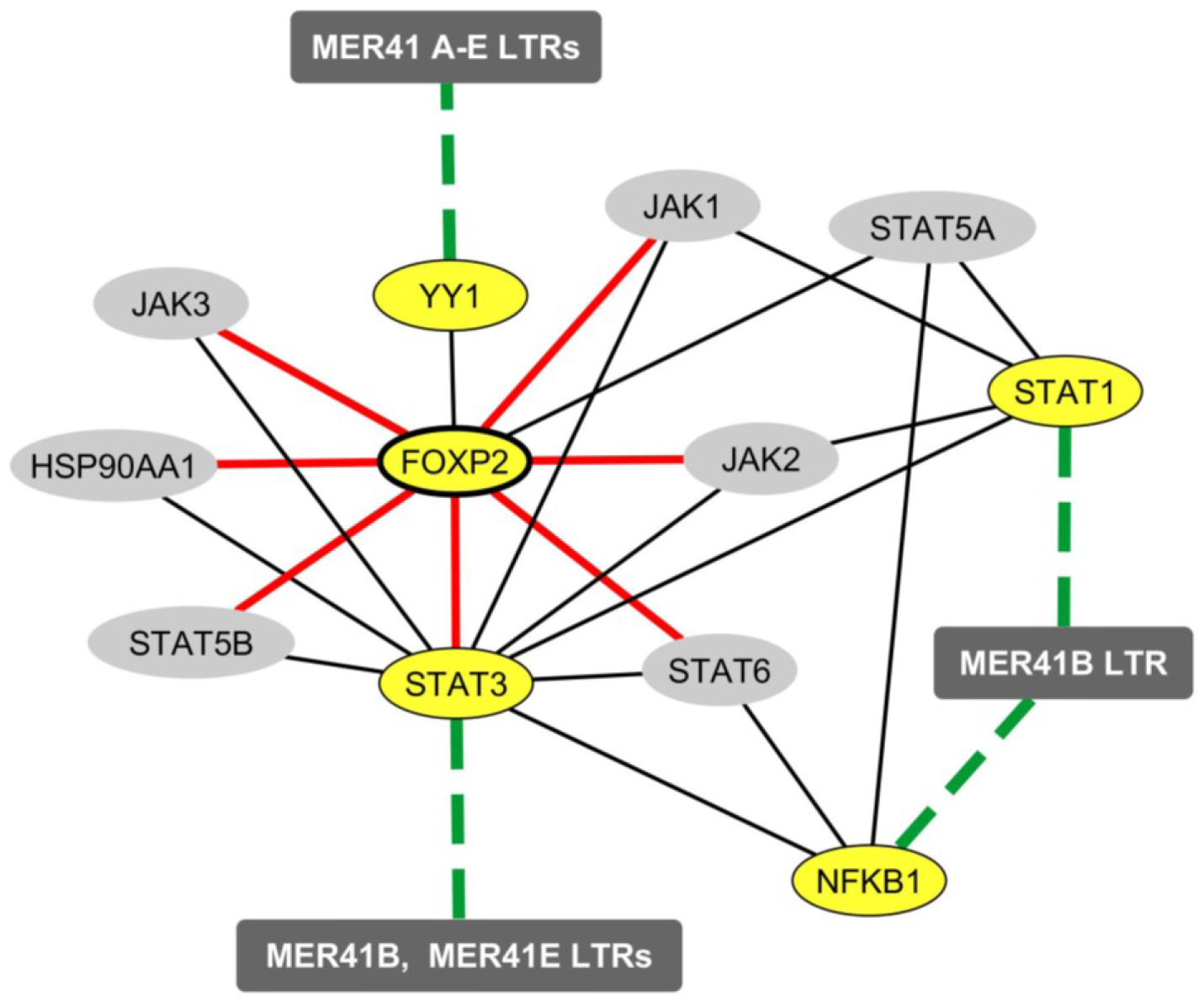
Mapping of the protein and functional network linking FOXP2 to immune TFs that bind MER41 LTRs in the promoter regions of cognition-related (ID-associated) genes. A survey of the human proteome was performed on the BioGRID database (Chatr-aryamontri et al., 2015) in order to map protein-protein interactions between FOXP2, STAT1, STAT3, NFKB1, YY1 and the molecules identified as being human-specifically regulated by FOXP2 in the neuronal cell line SH-SY-5Y (Oswald et al., 2017). Ellipse shapes highlighted in yellow represent FOXP2, its protein partner YY1 and the 3 immune TFs that bind MER41 LTRs in the promoter regions of cognition-related (ID-associated) genes (namely STAT1, STAT3 and NFKB1). Ellipse shapes highlighted in grey represent molecules that interact with these TFs and are putatively regulated by FOXP2 in a human-specific manner. Rectangles represent MER41 families displaying LTRs in the promoter regions of cognition-related (ID-associated) genes. Black lines indicate protein-protein interactions. Dashed green lines indicate protein-DNA interactions between TFs and MER41 LTRs located in the promoter regions of cognition-related (ID-associated) genes. Red lines indicate molecules that are targeted by FOXP2 in a human-specific manner

### 8 Human neural cells express key immune genes involved in the immune/MER41/cognition pathway

To further assess the relevance of our findings, we surveyed 2 independent databases which allows determining the neural expression of key genes putatively involved in immune/MER41/cognition pathway. These databases comprise: i) the “TISSUES” database (Palasca et al., 2018) which compiles manually curated expression results obtained in 4 distinct expression atlases (Clark et al., 2007; Fagerberg et al., 2014; Krupp et al., 2012; Su et al., 2004) covering a large range of normal human tissues and ii) the recently-launched “Brain RNA-Seq” database (Zhang et al., 2016) which allows exploring expression profiles observed in primary cultures of human neurons, astrocytes or macrophages/microglia. In our survey, the lymphocyte-specific gene markers *CD3G* and *ZAP70* were used as negative controls. Retrieved data showed that in cultured human neurons, *CD3G* and *ZAP70* are expressed at levels considered as bellow the detection threshold (data supplement 8). In contrast, all the immune genes examined putatively in the immune/MER41/cognition pathway (i.e *NFKB1*, *STAT1*, *STAT3*, *YY1*, *IL6R*, *IL6ST and IL6*) were found to be expressed at detectable levels in cultured human neurons (data supplement 8). The same results were retrieved when assessing mRNA levels in other neural cell types including fetal astrocytes, matures astrocytes, and macrophages/microglia (data supplement 8). Interestingly, in this expression database, *STAT1* mRNA levels reached higher levels in neurons than in macrophages/microglia (data supplement 8). Of potential interest also, retrieved data showed that IL-6 is constitutively expressed by cultured macrophages/microglia derived from the brains of humans but not mice (data not shown). Regarding the expression pattern of candidate genes in normal human tissues, we observed that, as expected, *CD3G* and *ZAP70* exhibited their higher levels of expression in lymphoid tissues (e.g. thymus, tonsils or lymph nodes) (data supplement 8). However, surprisingly, the prototypical immune-related genes *STAT1*, *STAT3*, *NFKB1*, *IL6R* and *IL6ST* were reported to display their highest (or second highest) levels of expression in the human brain (data supplement 8). Similar results were obtained for *YY1*. On another hand, high levels IL-6 were reported in the spinal cord and pons (data supplement 8). Altogether, these retrieved data indicate that all the immune players putatively involved in the immune/MER41/cognition pathway are actually expressed by human neural cells including neurons.

## DISCUSSION

We have found that, in the human genome, the promoter regions of ID-associated genes are uniquely enriched in MER41 LTRs. More specifically, 9 ID-associated genes that are putatively important in cognitive evolution exhibit MER41 LTRs in their promoter regions. As more than 100 families of HERV are integrated into our genome, it was important to determine whether our findings are specific to MER41 and to ID-associated genes, and if so to what extent. Among the 133 families of HERV explored here, MER41 is the only family whose LTRs were found with statistically high frequency in the promoter regions of ID-associated genes. It must be emphasized that, while many HERV families are inherited from ancestors common to all mammals, the MER41 family is detected exclusively in the genome of primates. Interestingly, we have observed substantial differences between humans and chimpanzees regarding the localization of MER41 LTRs in the promoter regions of ID-associated genes. Moreover, differences are also pointed out between humans and chimpanzees, regarding the protein sequences of 4 TFs (YY1, ESR1, SP1 and NANOG) binding MER41 LTRs, as well as 6 molecules (CDH15, CEP290, GAMT, DDHD2, GCSH and DEAF1) encoded by targeted cognition-related (ID-associated) genes. These results suggest that the MER41 family of HERVs could have been involved in cognitive changes after our split from chimps. In this scheme, infection and horizontal transmission of the exogenous virus from which MER41 HERVs derive, would have occurred in a community of primate ancestors and would have led to germline infection, followed by vertical transmission and, *in fine*, endogenization. If so, genomic evolution from these primate ancestors would have been, at least in part, dictated by the processes of HERV endogenisation and domestication, which is itself mainly dictated by the host’s immune system (Dewannieux and Heidmann, 2013). Accordingly, differences regarding the insertion sites of MER41 LTRs in the promoter region of a large range of genes, including cognition-related (ID-associated) genes, might have played roles in cognitive speciation. Conferring an evolutionary role to the immune/MER41/cognition pathway is further supported by the demonstration of human vs chimp differences at 3 levels of the immune/MER41/cognition pathway: i) TFs that bind MER41 LTRs, ii) insertions sites of MER41 LTRs and iii) cognition-related (ID-associated) genes with promoter-localized MER41 LTRs. It is worth noting that MER41 LTRs are not enriched in the promoter regions of ASD- or schizophrenia-related genes. This finding suggests that selected aspects of cognitive evolution in the primate genus are linked to MER41.

Our work also indicates that, in humans, immune regulation of the MER41/cognition pathway is not limited to IFNγ and its main downstream signaling molecule, STAT1. Indeed, the MER41 LTRs located in the promoter regions of cognition-related (ID-associated) genes harbor TFBSs for a group of 5 interacting immune-related TFs (STAT1, STAT3, NFKB1, YY1 and CEBPB) which are themselves functionally linked to multiple cytokines including IFNγ. Moreover, in this functional network, the prototypical pro-inflammatory cytokine IL-6 rather than IFNγ appears to be the main hub. Thus, cognitive evolution after our split from chimps might have been influenced by the process of endogenization and domestication of MER41 HERVs and by the parallel genomic evolution of immune genes. In this view, it is worth noting that, overall, immune genes harbor the highest levels of purifying selection in the human genome, which reflects the key functions of immunity in the defense against life-threatening infectious agents (Deschamps et al., 2016). This is notably the case for STAT1 (Deschamps et al., 2016).

Owing to its putatively important role in the immune/MER41/cognition pathway, YY1 deserves particular attention. Indeed, YY1 binding sites are observed in the LTRs from all MER41 subtypes (MER41 A to E) and YY1 is a direct protein partner of both NFKB1 and FOXP2, two TFs exerting major roles in immunity and language respectively. Moreover, YY1 is not only recognized as being crucially involved in CNS development (as notably shown in the inherited brain disorder “Gabriel-de Vries syndrome”), but also as exerting major functions in the immune system. In particular, YY1 was demonstrated to inhibit differentiation and function of regulatory T cells by blocking Foxp3 expression (Hwang et al., 2016) and to regulate effector cytokine gene expression and T(H)2 immune responses (Guo et al., 2008). We previously proposed that the nervous and immune systems have somehow co-evolved to the benefits of both systems, particularly regarding cognitive evolution, including our language-readiness (Benítez-Burraco and Uriagereka, 2016; Nataf, 2017a). In this context, YY1 may represent a new molecular connection between immunity and cognition and, even more specifically, between immunity and speech or language more generally.

NFKB1 may also draw specific interest since its neuronal expression was reported to be essential to behavior and cognition in both invertebrates and mammals (Dresselhaus et al., 2018; Kaltschmidt and Kaltschmidt, 2009; Mattson and Meffert, 2006; Meffert and Baltimore, 2005). In the central nervous system of rodents, components of the NFKB complex are detectable in neuronal processes and in synapses under physiological conditions (Dresselhaus et al., 2018; Salles et al., 2014). Moreover, synaptic transmission as well as exposure to neurotrophins activate the NFKB pathway in neurons (Kaltschmidt and Kaltschmidt, 2009; Mattson and Meffert, 2006; Meffert and Baltimore, 2005). In turn, NFKB activation in neurons triggers the transcription of multiple neuronal genes that may favor cognition and shape behavior. This is notably the case for neuropeptide Y and BDNF (Snow and Albensi, 2016).

The immune-mediated retrotranscription of specific HERVs was shown possibly to negatively influence the outcome of CNS disorders (Douville et al., 2011; Douville and Nath, 2017; Kremer et al., 2013; Küry et al., 2018). While our work unravels the putative evolutionary-determined advantage conferred by the immune/MER41/cognition pathway in human, it also points to the potential weaknesses that are inherent to such a pathway. Indeed, our data indicates that non-inherited forms of ID might result from alterations of the immune/MER41/cognition pathway. Such a dysfunction may be induced by the untimely or quantitatively inappropriate exposure of neurons to specific cytokines, notably IL-6 or IFNγ, which physiologically shape neurotransmission (Baier et al., 2009; Chourbaji et al., 2006; Gruol, 2015; Litteljohn et al., 2014; Victório et al., 2010) and are possibly involved in the immune/MER41/cognition pathway.

To conclude with a deliberately provocative view of cognitive evolution itself, a potential implication of our findings could be that cognitive evolution among primates might to some extent be a “bystander effect” of the immune system evolutionary adaptation to infections in general and, in particular, to retroviral infections. Excluding this hypothesis would lead to consider a related and barely less provocative view in which the last steps of our cognitive evolution might have promoted our immune defenses against human-specific infectious agents; for instance, evolving different languages as a sort of “protective barrier” against infectious diseases for the human groups speaking them. In any case, our work reinforces the notion of neuroimmune co-evolution that we previously put forward (Benítez-Burraco and Uriagereka, 2016; Nataf, 2017a). In this general frame, we would like to propose that, besides the potential role of endogenous immune cues (Nataf, 2017a, 2017b), immune signals triggered by infectious agents, might have been important to cognitive evolution. In particular, depending on their pathogenicity, such infectious agents could have exerted a neuroimmune selection pressure over millions of years (e.g., via the self-domestication of HERVs) or during short periods of time (e.g., via the occurrence of life-threatening epidemics of viral or bacterial infections). In this view, our findings provide general support to the hypothesis previously enunciated by Piattelli-Palmarini and Uriagereka (Piattelli-Palmarini and Uriagereka, 2004), updated by Benítez-Burraco and Uriagereka (Benítez-Burraco and Uriagereka, 2016) which states that the recent emergence of linguistic skills would have been triggered by a fast propagating virus.

## Acknowledgments

We thank Marine Guillen from Lyon-1 University (Histology Laboratory, UFR Lyon-Est) for the technical help she provided on internal quality control of bioinformatics analyses. The first version of this paper has been rendered available on the preprint platform bioRxv (Nataf et al., 2018) under the following doi: https://doi.org/10.1101/434209

## Authors contribution

SN performed the bioinformatics analyses and wrote the paper; ABB and JU co-wrote the paper.

