## Supplementary material for "The promoter regions of intellectual disability-associated genes are uniquely enriched in LTR sequences of the MER41 primate-specific endogenous retrovirus: an evolutionary connection between immunity and cognition": data supplement 5

**METHODS**

We queried the EnHERV data base and web tool to identify human genes harboring MER41 LTR sequence(s) in the promoter region located 2 KB upstream the TSS. A set of 79 coding genes was identified among which 9 are causally kinked to intellectual disability. For each of these 9 genes we checked the presence, nature and precise localization of MER41 LTR sequences in the promoter region of the Homo sapiens genome. To achieve this goal we used the UCSC genome browser (<https://genome.ucsc.edu/cgibin/hgGateway?hgsid=687072117_GjGeju9Eo1QozfesGAcSlA7ACB8O>) and explored the Human genome assembly GRCh38/hg38. We then compared these results with those obtained when exploring the Chimpanzee genome assembly CSAC 2.1.4/panTro4. Comparisons were performed regarding MER41 LTR sequences and localizations in the promoter regions of the 9 identified human genes and their orthologs in the Chimpanzee. The website LALIGN (<https://embnet.vital-it.ch/software/LALIGN_form.html>) was used to perform sequence comparisons.

**RESULTS**

For each gene analyzed are shown the position of the human gene, the position of the identified MER41 LTR in the promoter region, the position of the Chimpanzee ortholog gene, the position of the corresponding MER41 LTR sequence in the promoter region of the Chimpanzee ortholog gene and a sequence comparison between the MER41 LTR sequences observed in the Human vs Chimpanzee genomes. Differences are highlighted in green.

***CDH15***

Human gene Position: [chr16:89238163-89261900](https://genome.ucsc.edu/cgi-bin/hgTracks?hgsid=687054739_csNE3yGaHuMSAwfaXhh00zkTF8hA&db=hg19&position=chr16%3A89238163-89261900)

MER41A LTR Position: [chr16:89237361-89237425](https://genome.ucsc.edu/cgi-bin/hgTracks?hgsid=687054739_csNE3yGaHuMSAwfaXhh00zkTF8hA&db=hg19&position=chr16%3A89237361-89237425)

TATCAGAGGCGTGTGAACCAGAGCAGTTCCATACTGAATAAGAGCTGGGC

AAAATGAGGTCGCGA

Chimpanzee gene Position: [chr16:88903613-88918583](https://genome.ucsc.edu/cgi-bin/hgTracks?hgsid=669225851_pNAM0kIo2MU11Vr98yuzz1P8eGaU&db=panTro4&position=chr16%3A88903613-88918583)

**No MER41 LTR upstream TSS (up to 10 Kb) in the chimpanzee genome**

***CEP290 (reverse strand in Human and Chimpanzee)***

Human gene Position: [chr12:88049013-88142216](https://genome.ucsc.edu/cgi-bin/hgTracks?hgsid=669228021_j7aaTzSkI8RbokbSUd1VqBAymqaO&db=hg38&position=chr12%3A88049013-88142216)

MER41B LTR Position: chr12:88143723-88144293 TGTCAGAGGTATTTGAATCACAACAACTCCATCTTGAATAGGGGCTGGGT

AAAATAAGGGTGAGACCTACTGGGCTGCATTCTCAGGATGTCTGTCAGTC

ATTCTAAGTCACAGGATGAGATAGGAGGTGGGCACAAGATACTGATCATA

AAGACTTTGCTGATAAAACAGCATGCAGTAAAGAAGCCAGCCAAATCCCA

CCAAATCCAAGATGGCGAGAAAGGTGGGACGTCTGCTCGTTCTCACTGCT

CATTATACACTAATTATAATGCATTAGCATCCTAAAAGACACTCCCACCA

GTGCTATGACAGTTTACAGATGCCATGGCAATGTGAAGAAGTTACTCTAT

ATAGTCTAAAAAGGGGAGGAACACTCAGATCTAGGAATGGCCTACCCTCT

TCCCAGAAAACTCATGAATAACCCACCCCTTGTTTAGCATATAATCAAGA

AATAACTATAACAATCCTTAGTCTACCACCTCCAATGGGGGTGTATGCAC

CTCCAATTCTTTCTTGGCGAGATCCAGGAGCCCTCTCCTGGGGTCCAGAT

CAGAACACCTTTCCAGAAACA

Chimpanzee gene Position: [chr12:88199730-88292316](https://genome.ucsc.edu/cgi-bin/hgTracks?hgsid=669225851_pNAM0kIo2MU11Vr98yuzz1P8eGaU&db=panTro4&position=chr12%3A88199730-88292316)

MER41B LTR Position: chr12:88294737-88295307 TGTCAGAGGTATTTGAATCACAACAACTCCATCTTGAATAAGGGCTGGGT

AAAATAAGGGTGAGACCTACTGGGCTACATTCTCAGGATGTCTGTCAGTC

ATTCTAAGTCACAGGATGAGATAGGAGGTGGGCACAAGATACTGATCATA

AAGACTTTGCTGATAAAACAGCATGCAGTAAAGAAGCCAGCCAAATCCCA

CCAAATCCAAGATGGCAAGAAAGGTGGGATGTCTGCTCGTTCTCACTGCT

CATTATACACTAATTATAATGCATTAGCATCCTAAAAGACACTCCCACCA

GTGCTATGACAGTTTACAGATGCCATGGCAATGTGAAAAAGTTACTCTAT

ATGGTCTAAAGAGGGGAGGAACACTCAGATCTAGGAATGGCCTACCCTCT

TCCCAGAAAACTCATGAATAACCCACCCCTTGTTTAGCATATAATCAAGA

CATAACTATAACAATCCTTAGTCTACCACCTCCAATGGGGGTGTATGCAC

CTCCAATTCTTTCTTGGCGAGATCCAAGAGCCCTCTCCTGGGGTCCAGAT

CAGAACCCCTTTCCAGAAACA

Sequence comparison

MER41B LTR sequences 98.2% similar in 571 nt overlap (1-571:1-571)

**MER41B LTR sequence located beyond 2 Kb from TSS in the Chimpanzee genome**

***DDHD2***

Human gene Position: [chr8:38231491-38239092](https://genome.ucsc.edu/cgi-bin/hgTracks?hgsid=669326781_41Mn1ERTvQ5AbacfXL5WZCbta8YQ&db=hg38&position=chr8%3A38231491-38239092)

MER41B LTR Position chr8:38229500-38229570 GCTGCTTTGCCTATGGAGTAGCCATTCTTTTATTCCTTTACTTTCTTAAT AAACTTGCTTTCACTTTACTC

Chimpanzee gene position: [chr8:34772679-34805532](https://genome.ucsc.edu/cgi-bin/hgTracks?hgsid=669326781_41Mn1ERTvQ5AbacfXL5WZCbta8YQ&db=panTro4&position=chr8%3A34772679-34805532)

MER41B LTR Position: chr8:34769816-34769886 GCTGCTTTGCCTATGGAGTAGCCATTCTTTTATTCCTTTACTTTCTTAAT AAACTTGCTTTCACTTTACTC

Sequence comparison

MER41B LTR sequences 100% similar in 71 nt overlap (1-71:1-71)

**MER41B LTR sequence located beyond 2 Kb from TSS in the Chimpanzee genome**

***GCSH* (reverse strand in Human and Chimpanzee)**

Human gene Position: chr16:81,081,961-81,096,403

Mer41E LTR Position: [chr16:81096733-81097209](https://genome.ucsc.edu/cgi-bin/hgTracks?hgsid=669362729_4Bu6PfDdyatavfDk3XWR73CHc5Rw&db=hg38&position=chr16%3A81096733-81097209)

TGAGACAGGAACGTTACAGAGCAGTTGAAGGAGAATAGAAACTTCCAGGC

TGCAGTTCTGTCTAACAAAAGGAAACTGTTGAAATAGCTGCACAAGCTAT

GGGCTAAGACCCTGAAAAACCAGGGTGTGGGTCAACCTGGCTAAAACCAA

CTGGACCCAACGTGGTGTGGCTTTGACCTAGGCTTCACCTAGGACTCATT

AACATACTAAGTCACACACCCACCGGCACCACAACAATTCCGGGAACACC

CATATTTGGTGTAAAAATGGGTGGCACCACAGTTCTGACAAGTCTCCACC

TTATTCCAGAAACCTTTGTGTATATTCCATTTCTCAAAGAAACCCATAAA

GATGGAAACCCCAAGCCCCATTGTGTGGCCCTCTTTTGAGTCCGCCCTTT

CTTCAGTGTGTACTTTGCAGTAAACCTCTGTACTTTCACGACTTTCCGAC

TTGTTCTTGAATTCCTTCGTGAGGTGG

Mer41B LTR (1) Position: [chr16:81097577-81098100](https://genome.ucsc.edu/cgi-bin/hgTracks?hgsid=669362729_4Bu6PfDdyatavfDk3XWR73CHc5Rw&db=hg38&position=chr16%3A81097577-81098100) ACTTAGTCACAGGATGAGCTAGGGTTGGCACAAGATACAGGTCACAAAGT

CCCTGCTGATAAAATAGGATGCGGTAAAGGAGCTGGCCAAAACTCACCAA

AACCAAGATGGCGACCTCTGGTAGGTCTCACTGCTCATTATACGCTAATT

ATAATGCATTAGCATGTTAAAAAAAAAACAACTCCCACCAGCGTCATGAC

AGTTCACAAATGCAATGGCAACGTCCAGAGAAGTTACCCTATAGTCCAAA

AAGCGGAGGAACCGTTAGTTCTAGGAAATCCCTGCCCCTTTTCTGGAGAA

CTCTTGAATGATCCACCCCTTGTTTAGCATACAATCAAGAAATAACTATA

AGTACACTTAGTCGAGAAGCCATACTGCTGCTCTGTCTACGAAGAAGCCA

TTCTTTTTTTTCTTTACTTCTCTATTAAACGTGCTTTCACTTTATAGACT

CTCCCCAAATTCTTTCTTGCATGAGGTCCAAAAACCCTCTCTTGGGGTCT

GGATTGGGAGCCCTTTCTGTAACA

Mer41B LTR (2) Position: [chr16:81098360-81098449](https://genome.ucsc.edu/cgi-bin/hgTracks?hgsid=669362729_4Bu6PfDdyatavfDk3XWR73CHc5Rw&db=hg38&position=chr16%3A81098360-81098449)

GGTGTTTGAACAAGAACAACTCCATTTTGAATATGGGCTGGGTAAAATGA

GGCTGAGACTTGGTGGGCTGCATTCCCAGGAGGTTAGGCA

Chimpanzee gene Position: [chr16:80611909-80626701](https://genome.ucsc.edu/cgi-bin/hgTracks?hgsid=669362729_4Bu6PfDdyatavfDk3XWR73CHc5Rw&db=panTro4&position=chr16%3A80611909-80626701)
Mer41E Position: [chr16:80627068-80627524](https://genome.ucsc.edu/cgi-bin/hgTracks?hgsid=669362729_4Bu6PfDdyatavfDk3XWR73CHc5Rw&db=panTro4&position=chr16%3A80627068-80627524)

TGAGACAGGAACGATACAGAGCAGTTGAAGGAGAATAGAAACTTCCAGGC

TGCAGTTCTGTCTCACAAAAGGCAACTGTTGAAATAGCTGCACAAGCTAT

GGGCTAAGACCCTGAAAAACCAGGGTGTCGGTCAAGCTGGCTAAAACCAA

CTGGACCCAACGTGGTGTGGCTTCACCCGGGGACTCATTAACATACTAAG

TCACACACCCACCGGCACCACGACAATTCCGGGAACACCCATATTTGGTG

TAAAAATGGGTGGCACCACAGTTCTGAGAAGTCTCCACCTTATTCCAGAA

ACCTTTGTGCATATTCCATTTCTCAAAGAAACCCATAAAGATGGAAACCC

CAAGCCCCATTGTGTGGCCCTCTTTTGAGTCCGCCCTTTCTTCAGTGTGT

ACTTTGCAGTAAATCTCTGTACTTTCACGATTTTCCGACTTGTTCTTGAA

TTCCTTC

Sequence comparison

MER41E LTR sequences 94.9% similar in 468 nt overlap (1-468:1-457)

**No MER41B LTR (nor any MER41A, MER41C or MER41D LTR sequence) in the promoter region up to 10 Kb from TSS in the Chimpanzee genome**

***GAMT* (reverse strand in Human and Chimpanzee**)

Human gene Position: [chr19:1397026-1401570](https://genome.ucsc.edu/cgi-bin/hgTracks?hgsid=669895533_2EsU29stEejHan1BGHCNG2lANLjJ&db=hg38&position=chr19%3A1397026-1401570)

Mer41A LTR Position: [chr19:1402109-1402161](https://genome.ucsc.edu/cgi-bin/hgTracks?hgsid=687057765_ar4QVq85aYNB4qinwixFeOvIA01m&db=hg38&position=chr19%3A1402109-1402161)

GGCTGCTCTATGGAGTAGCCATTCATTTTTATTCCTTAATAAACTTGCTT

TCA

Chimpanzee gene Position: [chr19:1348385-1352645](https://genome.ucsc.edu/cgi-bin/hgTracks?hgsid=669225851_pNAM0kIo2MU11Vr98yuzz1P8eGaU&db=panTro4&position=chr19%3A1348385-1352645)

MER41A LTR Position: [chr19:1353292-1353348](https://genome.ucsc.edu/cgi-bin/hgTracks?hgsid=669225851_pNAM0kIo2MU11Vr98yuzz1P8eGaU&db=panTro4&position=chr19%3A1353292-1353348)

GGCTGCTCTGTCTATGGAGTAGCCATTCTTTTTTATTCCTTAATAAACTT

GCTTTCA

Sequence comparison

MER41A LTR sequences 91.2% similar in 57 nt overlap (1-53:1-57)

***BBS10* (reverse strand in Human and Chimpanzee)**

Human gene Position: [chr12:76344486-76348442](https://genome.ucsc.edu/cgi-bin/hgTracks?hgsid=669020285_Oo02M5JWFMD2pddc93VTU1Qr8Pnp&db=hg38&position=chr12%3A76344486-76348442)

Mer41A LTR Position: [chr12:76350292-76350834](https://genome.ucsc.edu/cgi-bin/hgTracks?hgsid=669020285_Oo02M5JWFMD2pddc93VTU1Qr8Pnp&db=hg38&position=chr12%3A76350292-76350834)

GTCAGAGGCATCTGAACCAGAGCAACTCCATCTTGAATAGGAGCTGGGTA AAATGAGGCTAAGATCTAGGCTGCATTCCCAGATGATTAAGGCATTCTAA GTCACAGGATGAGATAGGAGGTTGGCACAAGATACAGGTCATAAAGACCT TGCTGATAAAACAGGTTGCAGTAAAGAAGCCAGCCAAAACCCACCAAAAC CAAGATGGCCACGAGAGTGACCTCTGGTTGTCCTCACTGCTATACTCCCA CCAGCACCAAAGACAGTTTACAAATGCCATGGCAATGTCAGGAAGTTACC CTATATGGCCTAAAGAGGGGAGGCATAAATAATCCACTCCTTGTTTAGCA TATCATCAAGAAATAACCATAAAAATCGGCAACCAGCAGCCCTTGGGGCT GCTTTGCCTATGGAGTAACCATTCTTTTATTCCTCTTTCTTAATAAACTT GCTTTCACTTTATGGACTCGTCCTGAATTCTTTCTTGCATGAGATCCAAG AACCCTCTCTTGGGGTCTGGATTGGGACCCTTTTCCTCTAACA

Chimpanzee gene Position: [chr12:76265513-76269468](https://genome.ucsc.edu/cgi-bin/hgTracks?hgsid=669020285_Oo02M5JWFMD2pddc93VTU1Qr8Pnp&db=panTro4&position=chr12%3A76265513-76269468)

Mer41A LTR Position: [chr12:76271316-76271861](https://genome.ucsc.edu/cgi-bin/hgTracks?hgsid=669020285_Oo02M5JWFMD2pddc93VTU1Qr8Pnp&db=panTro4&position=chr12%3A76271316-76271861)

GTCAGAGGCATCTGAACCAGAGCAACTCCATCTTGAATAGGAGCTGGGTA AAATGAGGCTAAGATCTACTAGGCTGCATTCCCAGATGATTAAGGCATTC TAAGTCACAGGATGAGATAGGAGGTTGGCACAAGATACAGGTCATAAAGA CCTTGCTGATAAAACAGGTTGCAGTAAAGAAGCCAGCCAAAACCCACCAA AACCAAGATGGCCACGAGAGTGACCTCTGGTTGTCCTCACTGCTATACTC CCACCAGCACCAAAGACAGTTTACAAATGCCATGGCAATGTCAGGAAGTT ACCCTATATGGCCTAAAGAGGGGAGGCATAAATAATCCACTCCTTGTTTA GCATATCATCAAGAAATAACCATAAAAATCGGCAACCAGCAGCCCTTGGG GCTGCTTTGCCTATGGAGTAACCATTCTTTTATTCCTCTTTCTTAATAAA CTTGCTTTCACTTTATGGACTCGTCCTGAATTCTTTCTTGCATGAGATCC AAGAACCCTCTCTTGGGGTCTGGATTGGGACCCTTTTCCTCTGACA

Sequence comparison

MER41A LTR sequences 99.3% similar in 546 nt overlap (1-543:1-546)

***DEAF1* (reverse strand in Human and Chimpanzee)**

Human gene Position: [chr11:644220-695754](https://genome.ucsc.edu/cgi-bin/hgTracks?hgsid=669020285_Oo02M5JWFMD2pddc93VTU1Qr8Pnp&db=hg38&position=chr11%3A644220-695754)

Mer41A LTR Position: [chr11:696404-696448](https://genome.ucsc.edu/cgi-bin/hgTracks?hgsid=669020285_Oo02M5JWFMD2pddc93VTU1Qr8Pnp&db=hg38&position=chr11%3A696404-696448)

CTAGAGGCTGTCCTGCCTATGGAGTAGCCATTCTTTATTCCTTCA

Chimpanzee gene Position: [chr11:682184-745400](https://genome.ucsc.edu/cgi-bin/hgTracks?hgsid=669020285_Oo02M5JWFMD2pddc93VTU1Qr8Pnp&db=panTro4&position=chr11%3A682184-745400)

Mer41A LTR Position: [chr11:746064-746108](https://genome.ucsc.edu/cgi-bin/hgTracks?hgsid=669020285_Oo02M5JWFMD2pddc93VTU1Qr8Pnp&db=panTro4&position=chr11%3A746064-746108)

CTAGAGGCTGTCCTGCCTATGGAGTGGCCATTCTTTATTCCTTCA

Sequence comparison

MER41A LTR sequences 97.8% similar in 45 nt overlap (1-45:1-45)

***AP1S1***

Human gene Position: [chr7:101154405-101161276](https://genome.ucsc.edu/cgi-bin/hgTracks?hgsid=669020285_Oo02M5JWFMD2pddc93VTU1Qr8Pnp&db=hg38&position=chr7%3A101154405-101161276)

MER41D LTR (1) Position: [chr7:101153216-101153402](https://genome.ucsc.edu/cgi-bin/hgTracks?hgsid=669020285_Oo02M5JWFMD2pddc93VTU1Qr8Pnp&db=hg38&position=chr7%3A101153216-101153402)

GAAAAGAAAAACTTAAGACAGTGTGTTCCTCTGTTTGCTTTCTGAGGACG CCCTACTCCGTAAGGGTGTAGCTTTCAATAAACTCTCTCTTCTCATCGCA CTCTACGACTTGCTTTGGGTTCCTTCCTGCATGAGATCCAAGAATTCTCT CTTGGGGTCTGGATCGGGACCCCTTTTTCCAGTAACA

MER41D LTR (2) Position: [chr7:101152752-101152903](https://genome.ucsc.edu/cgi-bin/hgTracks?hgsid=669020285_Oo02M5JWFMD2pddc93VTU1Qr8Pnp&db=hg38&position=chr7%3A101152752-101152903)

AGTTTTGCTTTGATGTACTTACCTACTAGAATGTCAAGGATAGTTAAATT AACAGAATAATAAATTTTGTCATGCCATTGGCCCATCTGCACATTGACTC GGCTTAGTTTAGTCTCTACAAAAACAAGACCCCTATATAAGAAAAACTTA

AA

Chimpanzee gene Position: [chr7:101761797-101768633](https://genome.ucsc.edu/cgi-bin/hgTracks?hgsid=669020285_Oo02M5JWFMD2pddc93VTU1Qr8Pnp&db=panTro4&position=chr7%3A101761797-101768633)

MER41D LTR (1) Position: [chr7:101760616-101760802](https://genome.ucsc.edu/cgi-bin/hgTracks?hgsid=669020285_Oo02M5JWFMD2pddc93VTU1Qr8Pnp&db=panTro4&position=chr7%3A101760616-101760802)

GAAAAGAAAAACTTAAGACAGTGTGTTCCTCTGTTTGATTTCTGAGGACG CCCTACTCCGTAAGGGTGTAGCTTTCAATAAACTCTCTCTTCTCACCGCA CTCTGCGACTTGCTTTGGGTTCCTTCCTGCATGAGATCCAAGAATTCTCT CTTGGGGTCCGGATTGGGACCGCTTTTTCCAGTAACA

MER41D LTR (2) Position: [chr7:101760145-101760303](https://genome.ucsc.edu/cgi-bin/hgTracks?hgsid=669020285_Oo02M5JWFMD2pddc93VTU1Qr8Pnp&db=panTro4&position=chr7%3A101760145-101760303)
TAATTATAGTTTTGCTTTGATGTACTTACCTACTAGAATGTCAAGGATAG TTAAATTAACAGAATAATAAATTTTGTCATGCCATTGGCCCATCTGCACA TTGACTCGGCTTAGTTTAGTCTCTACAAAAACAAGACCCCTATATAAGAA

AAACTTAAA

Sequence comparison 1

MER41D LTR (1) sequences 96.8% similar in 187 nt overlap (1-187:1-187)

Sequence comparison 2

MER41D LTR (2) sequences 100.0% similar in 152 nt overlap (1-152:8-159)

***ST3GAL5 (reverse strand in Human and Chimpanzee)***

Human gene Position: [chr2:85839148-85889034](https://genome.ucsc.edu/cgi-bin/hgTracks?hgsid=669020285_Oo02M5JWFMD2pddc93VTU1Qr8Pnp&db=hg38&position=chr2%3A85839148-85889034)

MER41C LTR (1) Position: [chr2:85889906-85890028](https://genome.ucsc.edu/cgi-bin/hgTracks?hgsid=669020285_Oo02M5JWFMD2pddc93VTU1Qr8Pnp&db=hg38&position=chr2%3A85889906-85890028) ACCACTTTACTTTCTTAATAAACTTGCTTTTACTTTGCACTGTGGACTTA CTCTGAATTGTTTCTTGCGTGAGATCGAAGAACCCTCTCTTGGGGTCTAG

ATCGGGACCCCTTTCCTGTAACA

MER41C LTR (2) Position: [chr2:85890279-85890657](https://genome.ucsc.edu/cgi-bin/hgTracks?hgsid=669020285_Oo02M5JWFMD2pddc93VTU1Qr8Pnp&db=hg38&position=chr2%3A85890279-85890657)

TGTCAGAAGTGTTCAAACCAGAGCGCCTCCATTTTGAGTGAGGGCTAGAA AAATGAGGCTGGGACTTGCTAGGCTGCATTTCCTGAAAGCTAGGCATTCC TAGCCTCTAGGTGTTTATGGTTAAGAGAACAAATTAATAATGCTTACTAA ACAGACCCAGACTTGAGAGAGTCCAGATATCCCGATATCTGGAGAACAAA GGCATTCCTAATTTTGCTTTAAAAATAATAATATTGATTATTGCAAAATA TAGTAATTAGGGAAAATTAATCCTTTATCACAAACCCTTGTAGCAGAAGA CATCTGCCCATATATACAAGCATTACAAGCATTGTACCTTAGTTGGATAT GTCCCTCCTCTTATTTTCAGGAATGCCCT

Chimpanzee gene Position: [chr2A:87033968-87085089](https://genome.ucsc.edu/cgi-bin/hgTracks?hgsid=669020285_Oo02M5JWFMD2pddc93VTU1Qr8Pnp&db=panTro4&position=chr2A%3A87033968-87085089)

MER41C LTR (1) Position: [chr2A:87085961-87086083](https://genome.ucsc.edu/cgi-bin/hgTracks?hgsid=669020285_Oo02M5JWFMD2pddc93VTU1Qr8Pnp&db=panTro4&position=chr2A%3A87085961-87086083)

ACCACTTGACTTTCTTAATAAACTTGCTTTTACTTTGCACTGTGGACTTA CTCTGAATTGTTTCTTGCGTGAGATCGAAGAACCCTCTCTTGGGGTCTAG

ATCGGGACCCCTTTCCTGTAACA

MER41C LTR (2) Position: [chr2A:87086334-87086712](https://genome.ucsc.edu/cgi-bin/hgTracks?hgsid=669020285_Oo02M5JWFMD2pddc93VTU1Qr8Pnp&db=panTro4&position=chr2A%3A87086334-87086712)

TGTCAGAAGCGTTCAAACCAGAGCGCCTCCATTTTGAGTGAGGGCTAGAA AAATGAGGCTGGGACTTGCTAGGCTGCATTTCCTGAAAGCTAGGCATTCC TAGCCTCTAGGTGTTTATGGTTAAGGGAACAAATTAGTGATGCTTACTAA ACAGACCCAGACTTGAGAGAGTCCAGATATCCCGATATCTGGAGAACAAA GGCATTCCTAATTTTGCTTTAAAAATAATAATATTGATTATTGCAAAATA TAGTAATTAGGAAAAATTAATCCTTTATCACAAACCCTTGTAGCAGAAGA CATCTGCCCATATATACAAGCATTACAAGCATTGTACCTTAGTTGGATAT GTCCCTCCTCTTATTTTCAGGAATGCCCT

Sequence comparison 1

MER41C LTR (1) sequences 99.2% similar in 123 nt overlap (1-123:1-123)

Sequence comparison 2

MER41C LTR (2) sequences 98.7% similar in 379 nt overlap (1-379:1-379)
