## Supplementary material for "The promoter regions of intellectual disability-associated genes are uniquely enriched in LTR sequences of the MER41 primate-specific endogenous retrovirus: an evolutionary connection between immunity and cognition": data supplement 6

**METHODS**

We used the UniProt-provided web tool “Align” (<https://www.uniprot.org/align/>) to perform protein sequences comparisons between humans and chimpanzees. This site provides an easy way to identify amino acid dissimilarities and to determine whether or not such dissimilarities are located in sequences that are known or predicted to exert specific functions. When different isoforms of chimpanzee ortholog proteins were indexed, we took into account only the isoform displaying the highest % of homology with the reference human protein. Dissimilarities occurring in “coiled coil” sequences were not considered as functionally-relevant except when a specific function was assigned to such coiled coil regions.

**RESULTS**

For each protein analyzed, are shown the % of homology between humans and chimpanzees as well as, when applicable, the number of amino acid dissimilarities and their localization in a sequence with known or predicted function. When existing, dissimilarities leading to putative functional differences are highlighted in green. This analysis was performed for 2 sets of proteins: i) TFs that bind MER41 LTRs in the promoter regions of ID-associated genes, ii) proteins encoded by ID-associated genes exhibiting a promoter-localized MER41 LTR.

*TRANSCRIPTION FACTORS BINDING MER41 LTRs IN THE PROMOTER REGIONS OF CANDIDATE COGNITION-ASSOCIATED GENES*

**Human YY1: 99.7% homology with Pan troglodytes closest ortholog**

1 amino acid dissimilarity in the “Interaction with the SMAD1/SMAD4 complex” region

**Human ESR1: 99.1% homology with Pan troglodytes closest ortholog**

1 amino acid dissimilarity in the “Nuclear Receptor Ligand Binding Domain (NR-LBD)”

**Human NANOG: 98.3% homology with Pan troglodytes closest ortholog**

3 amino acid dissimilarities in the “Sufficient for strong transactivation activity” region

**Human SP1: 99.8% homology with Pan troglodytes closest ortholog SP1**

1 amino acid dissimilarity in the “repressor domain” region

**Human STAT3: 100% homology with Pan troglodytes closest ortholog**

**Human NFKB1: 100% homology with Pan troglodytes closest ortholog**

**Human STAT1: 100% homology with Pan troglodytes closest ortholog**

**Human BATF: 100% homology with Pan troglodytes closest ortholog**

**Human CEBPB: 100% homology with Pan troglodytes closest ortholog**

**Human CTCF: 100% homology with Pan troglodytes closest ortholog**

**Human EBF1: 100% homology with Pan troglodytes closest ortholog**

**Human FOSL1: 100% homology with Pan troglodytes closest ortholog**

**Human FOSL2: 100% homology with Pan troglodytes closest ortholog**

**Human GATA2: 100% homology with Pan troglodytes closest ortholog**

**Human JUN: 100% homology with Pan troglodytes closest ortholog**

**Human POU5F1: 100% homology with Pan troglodytes closest ortholog**

**Human SPI1: 100% homology with Pan troglodytes closest ortholog**

**Human USF1: 100% homology with Pan troglodytes closest ortholog**

**Human GATA4: 100% homology with Pan troglodytes closest ortholog**

**Human EGR1: 99.6% homology with Pan troglodytes closest ortholog**

No difference in sequences with known or predicted functions

**Human ELF1: 99.8% homology with Pan troglodytes closest ortholog**

No difference in sequences with known or predicted functions

**Human ELK4: 99% homology with Pan troglodytes closest ortholog**

No difference in sequences with known or predicted functions

**Human FOS: 99.7% homology with Pan troglodytes closest ortholog FOS**

No difference in sequences with known or predicted functions

**Human GATA1: 99.7% homology with Pan troglodytes closest ortholog**

No difference in sequences with known or predicted functions

**Human GATA6: 99.3% homology with Pan troglodytes closest ortholog**

No difference in sequences with known or predicted functions

**Human JUNB: 99.7% homology with Pan troglodytes closest ortholog**

No difference in sequences with known or predicted functions

**Human JUND: 99.7% homology with Pan troglodytes closest ortholog**

No difference in sequences with known or predicted functions

**Human MEF2A: 98.2% homology with Pan troglodytes closest ortholog**

No difference in sequences with known or predicted functions

**Human NFE2: 99.7% homology with Pan troglodytes closest ortholog**

No difference in sequences with known or predicted functions

**Human POU2F2: 88.5% homology with Pan troglodytes closest ortholog POU2F2**

No difference in sequences with known or predicted functions

**Human SRF: 99.6% homology with Pan troglodytes closest ortholog**

No difference in sequences with known or predicted functions

**Human TAL1: 99.6% homology with Pan troglodytes closest ortholog**

No difference in sequences with known or predicted functions

***ID-ASSOCIATED GENES HARBORING A PROMOTER-LOCALIZED MER41 LTR***

**Human CDH15: 96% homology with Pan troglodytes closest ortholog**

5 amino acid dissimilarities in the “signal peptide” sequence

1 amino acid dissimilarity in the “cadherin 3” domain

1 amino acid dissimilarity in the “cadherin 4” domain

3 amino acid dissimilarities in the “cadherin 5” domain

**Human CEP290: 98.1% homology with Pan troglodytes closest ortholog**

35 amino acid dissimilarities in the “Self-association (with itself or C-terminus)” region

3 amino acid dissimilarities in the “Self-association (with itself or N-terminus)” region

**Human GAMT: 99.1% homology with Pan troglodytes closest ortholog**

2 amino acid dissimilarities in the “RMT2” (arginine N- methyltransferase 2-like) domain

**Human DDHD2: 99.8% homology with Pan troglodytes closest ortholog**

1 amino acid dissimilarity in the “DDHD” domain

**Human GCSH: 98.8% homology with Pan troglodytes closest ortholog**

2 amino acid dissimilarities in the “transit peptide” toward mitochondria sequence

**Human DEAF1: 99.1% homology with Pan troglodytes closest ortholog**

1 amino acid dissimilarity in the “interaction with LMO4” region

1 amino acid dissimilarity in the “MYND-type” zinc finger sequence

**Human ST3GAL5: 98.8% homology with Pan troglodytes closest ortholog**

No difference in sequences with known or predicted functions

**Human BBS10: 99.4% homology with Pan troglodytes closest ortholog**

No difference in sequences with known or predicted functions

**Human AP1S1: 100% homology with Pan troglodytes closest ortholog**
